# Force variability is a potential biomarker of motor impairment in hemispheric stroke survivors

**DOI:** 10.1101/2024.09.08.611881

**Authors:** Fandi Shi, William Zev Rymer, Jongsang Son

## Abstract

During voluntary isometric contractions of upper extremity muscles in individuals with chronic stroke, the magnitude of force variability appears to increase consistently as force increases. However, research on how such force variability changes with increasing motor impairment remains limited. This study aims to determine whether force variability is increased on the paretic side during either index finger abduction or elbow flexion in the same group of stroke survivors, and whether these changes are consistent across different muscles. Force variability was assessed using the standard deviation of force during sustained isometric contractions. Linear mixed-effects models were implemented to test whether force variability is changed on the paretic side post stroke, and whether such alterations show dependence on force level and on the degree of impairment. The results demonstrated a significant increase in force variability on the paretic side across force levels during finger abduction, while force variability for elbow flexion was increased only at high force levels. In addition, the force variability appears to increase as isometric elbow flexion force increases, whereas no clear trend was found during index finger abduction. The increase in force variability demonstrated moderate-strong dependence on the reduction in maximum muscle strength on the paretic side during elbow flexion, suggesting that monitoring force variability could potentially serve as a quantitative diagnostic tool for assessing severity of impairment in motor control and for raising potential mechanisms at the motor unit level.

## I. INTRODUCTION

Fluctuations in force (or torque) output within the low-frequency band (< 2 Hz) have been widely reported during voluntary sustained isometric contractions in intact human skeletal muscles [1] [2]. These variations are often quantified using the standard deviation (SD) or coefficient of variation (CoV) for force [2] [3]. While there have been some studies conducted on force variability in various populations, the mechanism underlying its generation remains unclear. Using advanced decomposition techniques, most studies with recordings of concurrently active motor units (MUs) have shown that high-threshold motor units are being activated at discharge rates significantly deviating from the optimal fusion frequency of their fast-twitch fibers, which are characterized by a short half-relaxation time [4] [5]. This mismatch between the firing rate and mechanical response of MUs may lead to larger force ripples, thus increased force variability observed at high contraction levels when high-threshold MUs are typically recruited. Expanding on the implication of this hypothetical condition, individuals with neurological disorders such as stroke, where MUs firing rates are reduced on the paretic side compared to normal [6], may experience greater force variability in paretic muscles. Although some studies have noted the increased CoV for force in hip and knee extensions post stroke [7] [8], there are limited evidence of the potential changes in force variability in upper limb muscles (i.e., arm and hand muscles) [9] [10] which typically exhibit more weakness following stroke. In this study, we aim to investigate and compare the changes in force variability generated in upper extremity muscles post stroke. These efforts will serve as a critical step towards the long-term goal of developing a simple biomarker to monitor motor function in clinical populations.

After stroke, various impairments have been documented, such as altered muscle activations, abnormal muscle synergies, more muscle fatigue, and increased muscle spasticity [11] [12] [13] [14] [15] [16]. These impairments adversely affect motor function and might compromise the ability to maintain precise force output in paretic muscles. Furthermore, impairments in muscle activation exhibit differences across different upper limb muscles following a stroke. There is a classical perception that motor recovery follows a proximal-to-distal gradient, implying more severe impairments associated with the hand intrinsic muscles [17]. Moreover, the modulation of arm and hand muscle activity by the nervous system involves different neural pathways. To be precise, the proximal arm motor neurons are more extensively modulated by interneurons within the cervical spinal cord, whereas hand muscles receive more direct input from corticospinal tract neurons [18] [19]. Taken together, the disruption to the corticospinal tract might disproportionately affect motor function involving hand muscles compared with the proximal arm post-stroke. Other features among different muscles, such as muscle architecture, muscle fiber type distribution, and recruitment/firing properties of MUs [20] [21] [22] [23] [24] could also contribute to different force generation and control ability. To date, limited study has examined whether the degree of impairment in force output is similar among different muscle groups. Such investigation could help assess the potential generalization of changes in force variability following a stroke or, if different changes are found, offer insights into the factors regulating force variability in different muscles.

Accordingly, the objective of the present study is to investigate whether force variability is increased on the paretic side during index finger abduction and elbow flexion in the same group of stroke survivors, involving first dorsal interosseous (FDI) and elbow flexors, respectively; and whether such reductions are uniform between muscles across the subjects. Considering alterations in MUs firing patterns after stroke as well as the potential differences in the control mechanisms and neuromechanical properties among muscles, we hypothesize that force variability is increased on the paretic sides post-stroke, and that such an increase differs between proximal (i.e., elbow flexors) and distal (i.e., FDI) muscles.

## II Methods

### A. Participants

Out of seventeen chronic hemiparetic stroke survivors recruited, twelve were included in this study due to their ability to perform both index finger abduction and elbow flexion. The demographic information of these subjects is described in Table 1. Before testing, informed consent was obtained from each participant. All procedures were approved by the Institutional review board at Northwestern University and complied with the Helsinki declaration compliance.

**TABLE 1.** Subject Demographic Information.

| Subject ID | Gender | Age (yrs) | Stroke Type | Year(s) Since Stroke | Paretic Side | MVIC Ratio (FA) | MVIC Ratio (EF) |
| --- | --- | --- | --- | --- | --- | --- | --- |
| 01 | M | 68 | I | 19 | R | 0.67 | 0.56 |
| 02 | M | 67 | I | 13 | L | 0.75 | 0.60 |
| 03 | M | 71 | I | 5 | L | 0.77 | 0.66 |
| 04 | M | 59 | I | 16 | R | 0.62 | 0.60 |
| 05 | M | 45 | H | 11 | R | 0.79 | 0.78 |
| 06 | M | 65 | H | 13 | R | 1.03 | 1.08 |
| 07 | F | 19 | I | 2 | L | 0.48 | 0.56 |
| 08 | F | 60 | I | 11 | R | 0.99 | 0.74 |
| 09 | F | 45 | I | 29 | R | 0.65 | 0.79 |
| 10 | F | 48 | I | 1 | L | 0.82 | 0.76 |
| 11 | M | 70 | H | 8 | L | 1.01 | 1.05 |
| 12 | F | 58 | H | 14 | R | 0.73 | 0.75 |

Note: 1) Gender: M–Male; F–Female. 2) Stroke type: I–Ischemic; H–Hemorrhagic. 3) Paretic side: R–Right; L–Left. 4) Maximum voluntary isometric contraction (MVIC) Ratio: MVIC on paretic side / contralateral side. 5) Muscle type: FA–index finger abduction; EF–elbow flexion.

Inclusion criteria were: 1) More than 6 months post a hemiparetic stroke; 2) Medically stable with medical clearance to participate; 3) Ability to provide informed consent; 4) No musculoskeletal conditions/surgeries/injuries in upper limb muscles; 5) Not currently taking any anti-spasticity medications; and 6) No cognitive impairment which could limit the comprehension of the tasks during testing.

### B. Experimental setup

During finger abduction, subjects were seated upright in a Biodex chair (Biodex Medical Systems, Shirley, NY, USA), with their upper arm resting on a support. The forearm was immobilized using a cast and placed within a ring interface attached to the platform, which was securely mounted to a steel table via four magnetic holders. To minimize potential mechanical interference from other hand muscles, the little, ring, and middle fingers were wrapped and strapped to a standing support. The thumb was extended and secured at approximately 45*º* relative to the index finger (Fig. 1a). Abduction force (Fx) was recorded from the proximal phalanx of the index finger, which was fixed to a ring-mount interface attached to a six degrees-of-freedom load cell (Gamma, ATI Inc., Apex, NC, USA; Resolution: 0.006 N).

**Fig. 1.**
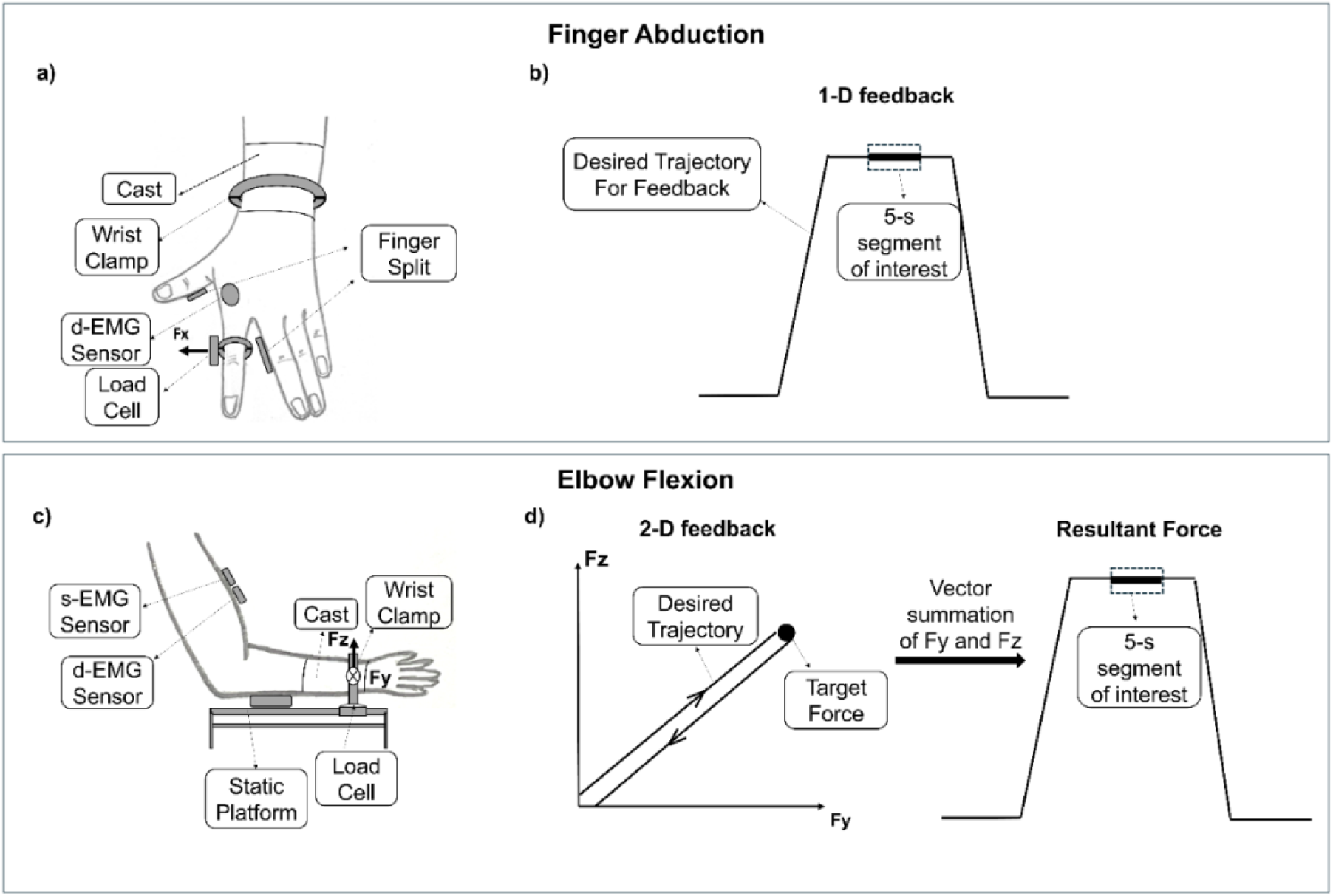
Experimental setup and protocol for visual feedback. Upper panel–during finger abduction: a) Experimental setup; b) 1-D force feedback and illustration for signal of interest. Lower panel–during elbow flexion: c) Experimental setup; d) 2-D force feedback and illustration for the resultant force including the signal of interest. s-EMG represents single differential electromyogram sensor, and d-EMG wireless Trigno Galileo sensor (Delsys Inc., Natick, MA, USA). Note that EMG signals were collected during the experiments, but not presented in the current manuscript.

During elbow flexion torque, participants were seated in an upright position with elbow flexed at 120*º*, shoulder abduction at 45*º*, shoulder flexion at 20*º*, and forearm pronation at 45*º*. The forearm was cast and immobilized in a ring. The fixture was attached with a six-degrees-of-freedom load cell (Delta, ATI Inc., Apex, NC, USA; Resolution: 0.06 N) to record voluntary forces generated at the wrist (Fig. 1c). Forces generated in the direction of Fy and Fz were used for further analyses. Data were collected at a sampling frequency of 2 kHz (NI USB-6259 BNC, National Instrument, Austin, TX, USA).

### C. Experimental protocol

For each subject, data collection was completed in two separate visits (one visit for one side, and paretic side first) for each of index finger abduction and elbow flexion protocols. For the first visit with the paretic side, participants were guided to perform maximum voluntary isometric contractions (MVICs) for at least 5 s, repeating the process three times with a 3-min rest interval between trials to minimize muscle fatigue. The average of the maximum values from each MVIC trial was used to determine the MVIC value for each subject. To provide a fair comparison between the sides at matched forces, the MVIC value from the paretic side was used to calibrate the target force level during submaximal contractions for both sides. The MVIC value on the contralateral side was also determined so that the MVIC ratio (i.e., MVIC on the paretic side / MVIC on the contralateral side; see Table 1) was calculated as an estimate of relative muscle weakness. Each subject was guided to perform three submaximal contractions at five contraction levels (i.e., 20–60%MVIC with an interval of 10%MVIC) in a random order. A 60-s break was provided between trials to minimize muscle fatigue, and additional breaks were provided upon request.

During index finger abduction, one-dimensional feedback of the abduction force was provided using a trapezoid trajectory display (Fig. 1b). The trajectory contains five segments: 1) 15-s quiescent baseline; 2) a ramp-up period with the moving rate at 10%MVIC/s; 3) a constant force plateau set at the prescribed target level for 12 s; 4) a ramp-down trajectory with the decreasing rate at 10%MVIC/s; and 5) 15-s quiescent period. During the plateau phase, participants were encouraged to maintain the contraction at the target level within ±3%MVIC. During elbow flexion, two-dimensional feedback was provided to track and maintain the same force for both dimensions (Fy and Fz) using a similar trapezoid trajectory. The resultant elbow flexion force was calculated as the vector summation of Fy and Fz (Fig. 1d).

### D. Data Analysis

Force signals were lowpass filtered using a 4th order Butterworth filter with a cutoff frequency of 10 Hz. A 5-s segment in the middle of the plateau was selected with minimum standard deviation. To eliminate the potential effects from extremely low-frequency motion artifact, a high-pass filter with the cutoff frequency of 0.2 Hz was applied. The standard deviation for force corresponding to the 5-s segment force was calculated to estimate force variability (hereafter called FSD). Fig. 2 illustrates typical force fluctuations over a 5-s interval at 40%MVIC on both sides during isometric index finger abduction (Fig. 2a) and elbow flexion (Fig. 2b) from a representative subject. The average value among three repetitions was calculated to determine FSD at each contraction level on each side within one subject. For each subject, these values from both sides (i.e., average FSD on the paretic side, called *FSD*_*Pare*_ ; and average FSD on the contralateral side, called *FSD*_*Contra*_) were then used to quantify the relative changes in force variability post stroke at each contraction level, called *FSD*_*Norm*_, using the following equation:

**Fig. 2.**
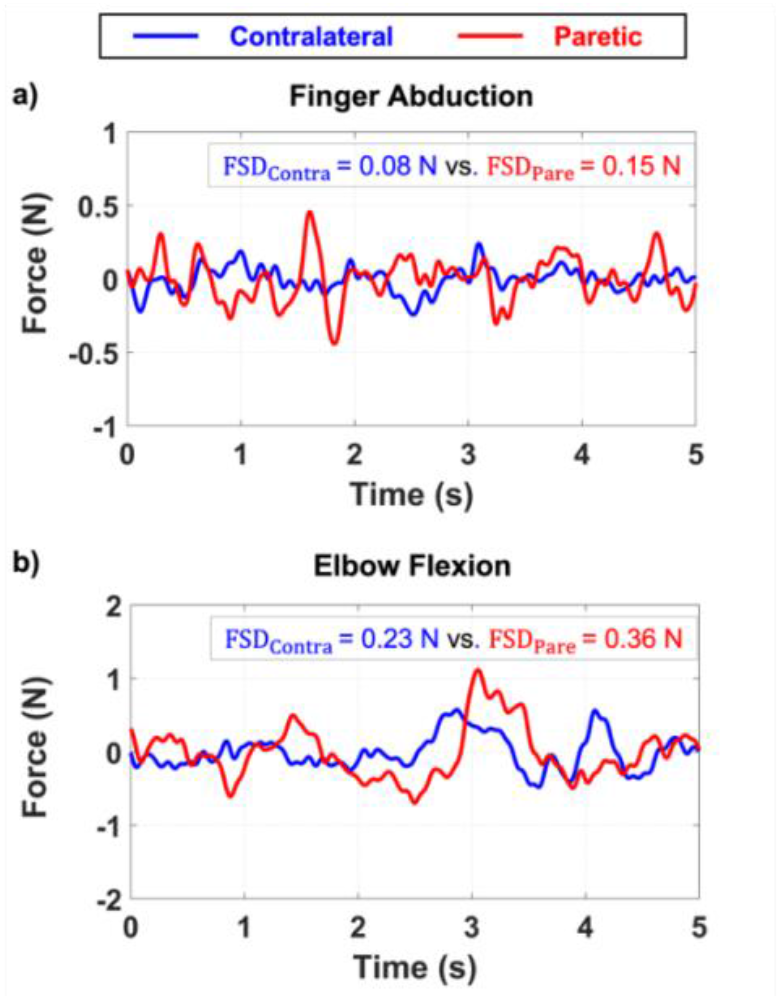
Typical force trials on both sides at 40%MVIC for 5 s during index finger abduction (a) and elbow flexion (b) from a representative subject.

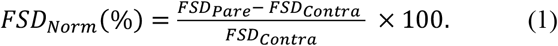

To understand the heterogeneity in muscle weakness and force variability, respectively, between the two muscle groups, we compared the MVIC ratio and FSD ratio (i.e., *FSD*_*Pare*_/*FSD*_*Contra*_) for the two muscles for each participant. The FSD ratio in each subject was determined as an average value of the FSD ratios across different contraction levels. For the sake of readability (i.e., the more the impairment on the paretic side, the smaller the MVIC ratio whereas the greater the FSD ratio), two comparison metrics were calculated as follows:

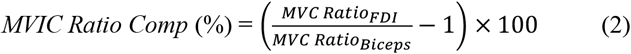

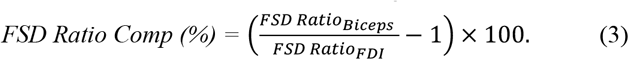

For instance, if the paretic FDI muscles are more affected in both the MVIC and force variability compared to the elbow flexors, both the comparison metrics are smaller than 0, and vice versa.

A fast Fourier transform was conducted to convert force signals from the time domain into the frequency domain. For each subject, the magnitude across all bins within 10 Hz of each trial was normalized by the maximum magnitude found on the paretic side at 60%MVIC, which was calculated by the average of the repeated trials at this force level. For each subject, the average of all the normalized magnitude spectrums across the repeated trials at each contraction level for each muscle was calculated for further analyses.

### E. Statistical analysis

All the statistical analyses were done using MATLAB (R2021b, The MathWorks, Inc., Natick, MA, USA). Any significance was determined when the p-value was less than 0.05.Data was shown as Mean ± standard error of mean (SEM) in main text and figures. The Kolmogorov-Smirnov test was conducted to assess the normality of the MVIC values for both sides during both index finger abduction and elbow flexion. As the normality tests rejected the null hypothesis, a non-parametric paired t test was used to compare the MVIC values between the two sides during each task.

Linear mixed-effects models were implemented separately for each muscle, in order to mainly test 1) whether force variability is different between sides; and 2) whether the relative changes in force variability are different between muscles. For testing the first hypothesis, intercept, side, contraction level and interaction between side and contraction level were all set as fixed effects. For testing the second hypothesis, intercept and contraction level were treated as fixed effects. Subjects were treated as random effects for both tests. If any significance was found (p < 0.05), post-hoc analyses were performed, followed by multiple comparison adjustments using false discovery rate [25].

Statistical parametric mapping [26] is used to compare the differences in the force spectrum at each frequency bin between two sides during both finger abduction and elbow flexion.

The Spearman correlation analysis was performed to characterize the relation between the increase in force variability (quantified by *FSD*_*Norm*_) and muscle weakness (quantified by MVIC Ratio) during finger abduction and elbow flexion, respectively.

## III. Results

### A. Reduced strength on the paretic side

On average, the maximum strength generated on the paretic side was significantly reduced compared to the contralateral side during both index finger abduction (26.0 ± 2.6 N vs. 34.1 ± 3.5 N; t(11) = –4.077, p = 0.005) and elbow flexion (99.9 ± 9.5 N vs. 138.3 ± 13.5 N; t(11) = –5.033, p = 0.002), leading to an MVIC ratio less than 1 in 10 out of 12 stroke survivors (Table 1). Compared to the contralateral side, the average MVIC value on the paretic side reduced by ∼23% during index finger abduction and by ∼26% during elbow flexion.

### B. Changes in force variability between paretic and contralateral sides

During index finger abduction, FSD was a function of contraction level (F_4,104_ = 4.392, *p* = 0.003) and was also significantly greater on the paretic side compared to the contralateral side (F_1,104_ = 4.219, *p* = 0.042). Moreover, there was a significant interaction between contraction level and side (F_4,104_ = 2.900, *p* = 0.025). Post-hoc analysis revealed that FSD is greater on the paretic side at all the contraction levels (*p* < 0.05), except for at 20%MVIC (*p* = 0.057; Fig. 3a).

**Fig. 3.**
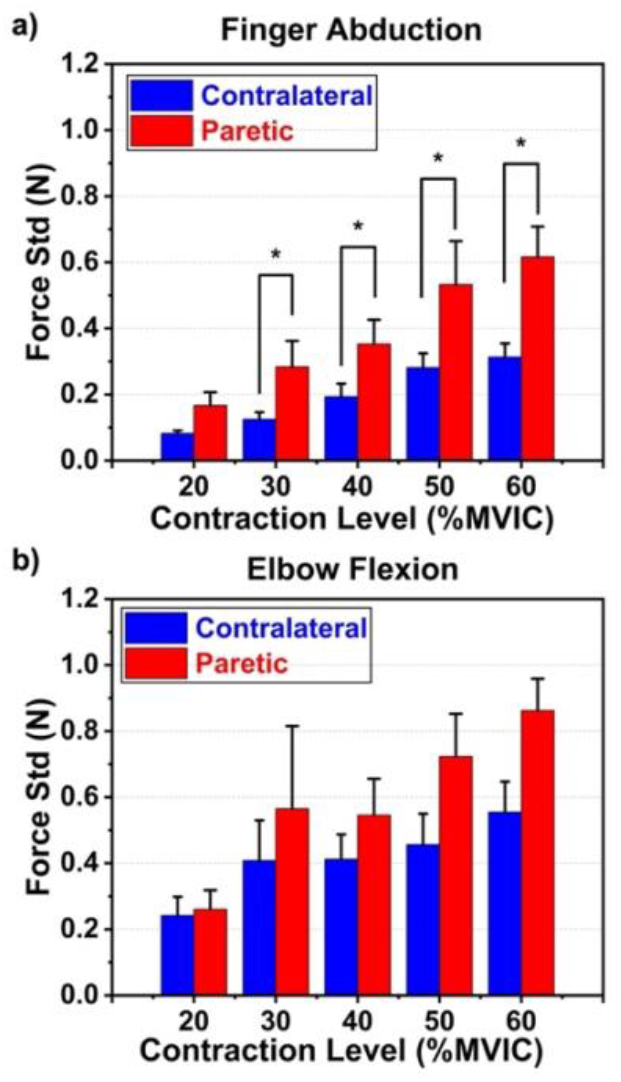
Summary results for force variability across all contraction levels on both sides during index finger abduction (a) and elbow flexion (b), averaged among all the subjects. Error bar indicates standard error of mean. Asterisk indicates a significant difference (\**p* < 0.05).

During elbow flexion, FSD was significantly increased with contraction level (F_4,100_ = 5.448, *p* = 0.001) but not significantly different by side (F_1,100_ < 0.0001, *p* = 0.993). In addition, the interaction between side and contraction level was significant (F_4,100_ = 3.656, *p* = 0.008), with greater disparity between two sides at high contraction levels (Fig. 3b). The supplementary Fig. 1 displays force variability against contraction levels on both sides for each participant.

### C. Dependence of changes in force variability on muscle selection

During finger abduction, the relative changes in force variability appear to be consistent at all the contraction levels (F_4,52_ = 1.649, *p* = 0.176; Fig. 4a). However, during the elbow flexion, there was an increasing trend in the relative changes in FSD magnitude as contraction level increased (F_4,50_ = 4.379, *p*

**Fig. 4.**
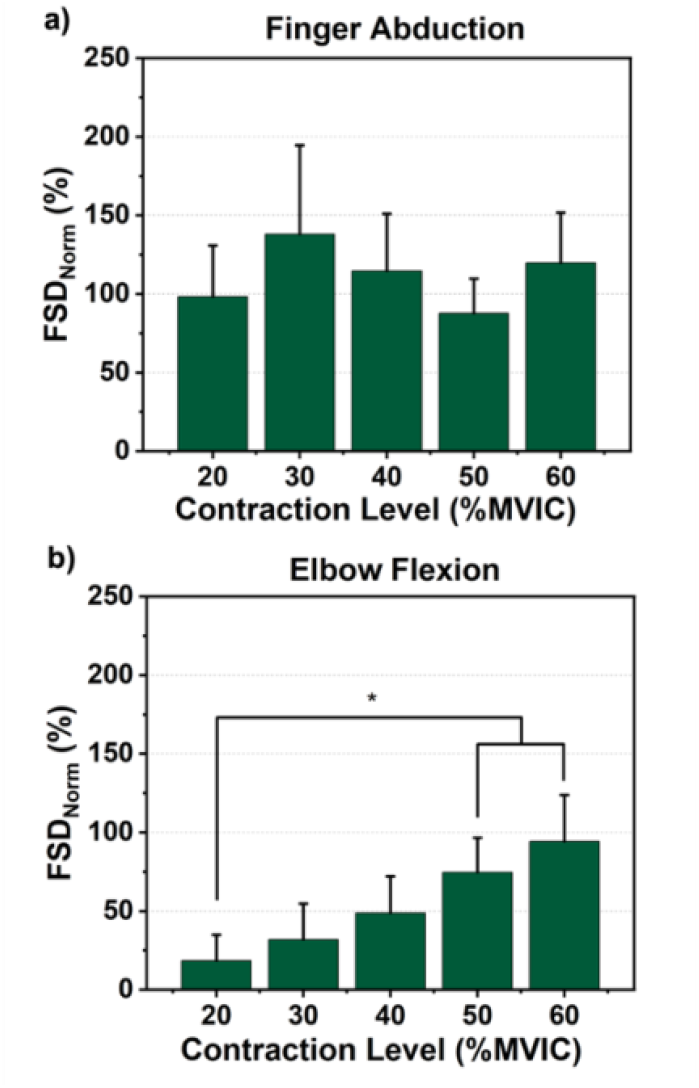
Relative changes in force variability between two sides at each contraction level during index finger abduction (a) and elbow flexion (b), averaged among all the subjects. Error bar indicates standard error of mean. Asterisk indicates a significant difference (\**p* < 0.05).

= 0.004), with significantly larger values at 50 and 60%MVIC compared to 20%MVIC (Fig. 4b).

### D. Low-frequency oscillations in force output

A dominant frequency lower than 2 Hz was clearly observed in force output on both sides during both index finger abduction and elbow flexion, as shown in the representative plots at 30%MVIC from an exemplar subject (Fig. 5). A secondary peak around 2 Hz was observed on the paretic side only during index finger abduction.

**Fig. 5.**
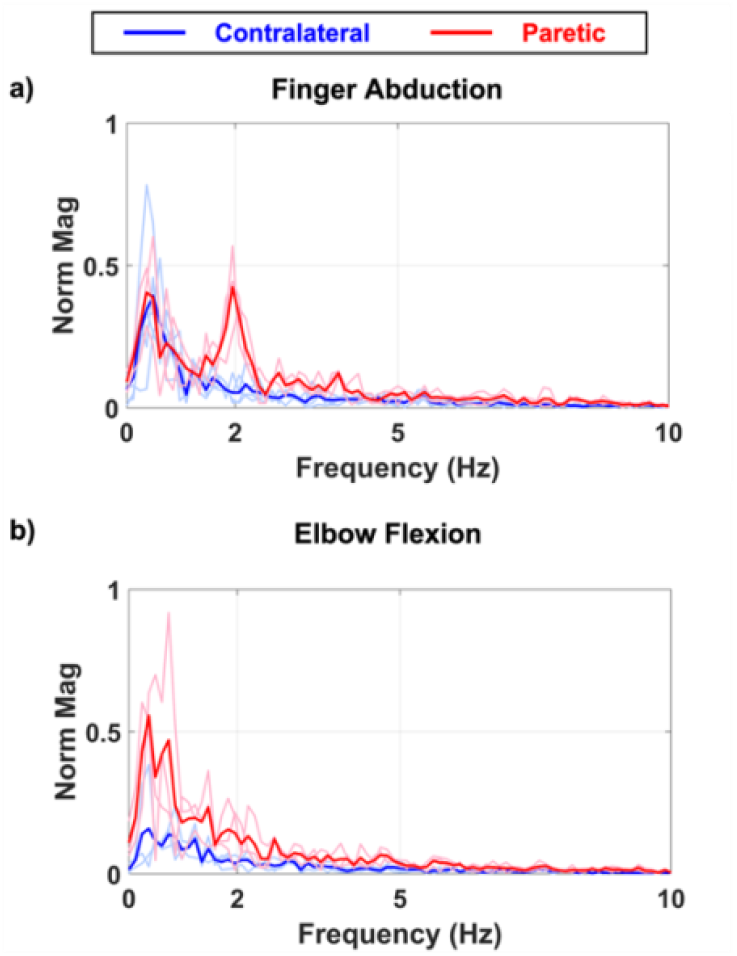
Representative results of the normalized magnitude distribution of force output at 30%MVIC on both sides during index finger abduction (a) and elbow flexion (b) from an exemplar subject. Note the remarkable secondary peak on the paretic side only during index finger abduction (dark red arrows). Norm Mag indicates the magnitude of the force spectrum normalized to the maximum magnitude of the force spectrum estimated at 60%MVIC. Light lines represent the individual trials and dark lines the average of the repetitions.

The peak frequency lower than 2 Hz was consistently observed across the force levels on both sides during index finger abduction and elbow flexion among the subjects (Fig. 6). For the comparison between sides, there was a trend for a secondary peak between 2 Hz and 5 Hz observable on the paretic side only during index finger abduction, with the evidence that a meaningful difference can be detectable at 30 and 50%MVIC compared to the contralateral side (*p* < 0.05), and at 40 and 60%MVIC (*p* < 0.1). However, such differences were not detected at higher frequencies beyond 2 Hz during elbow flexion.

**Fig. 6.**
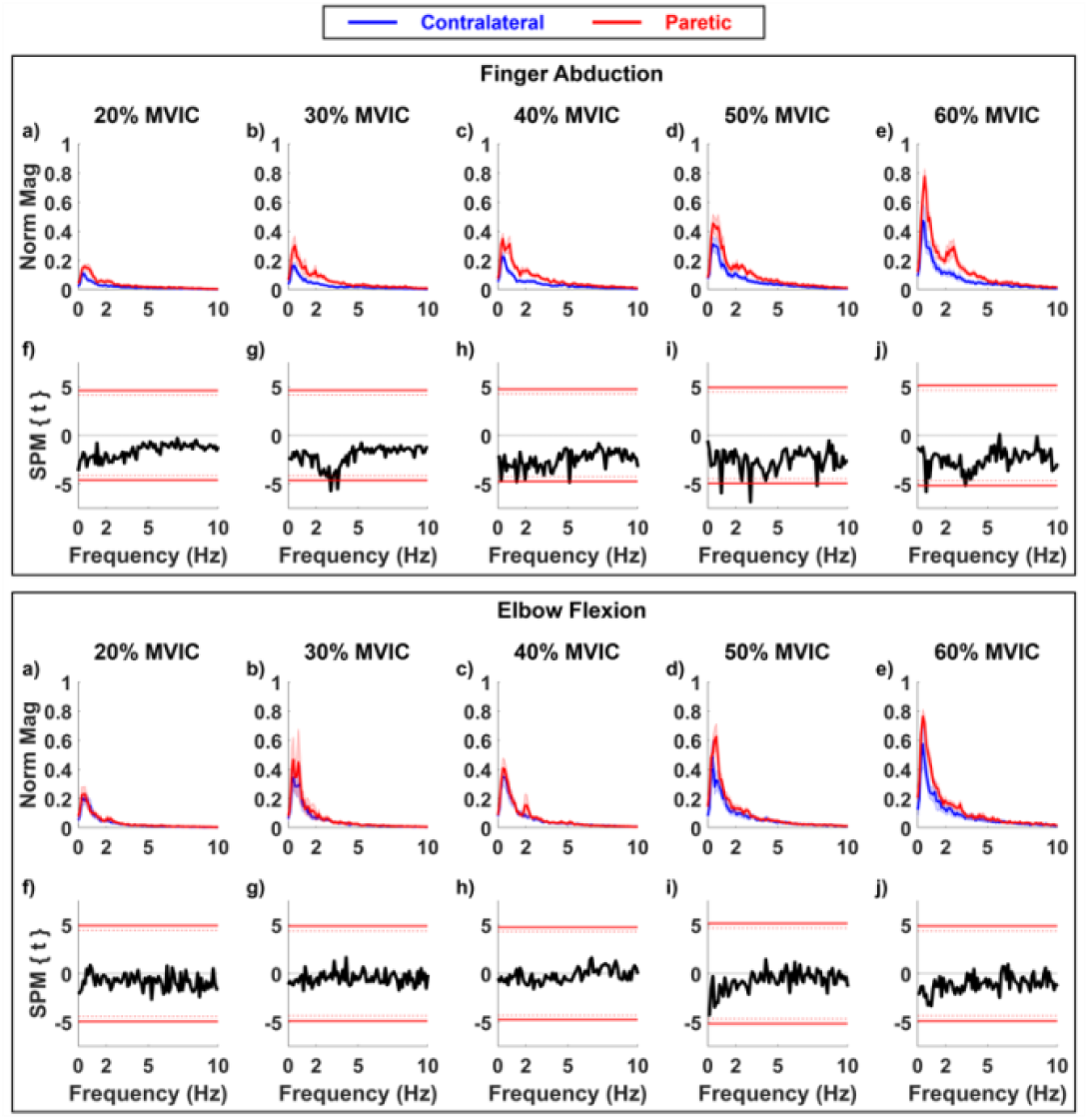
Normalized magnitude distribution of force output at each contraction level on both sides during index finger abduction (a–e) in the upper panel and elbow flexion (a–e) in the lower panel. The solid thick red lines represent the average among all the subjects, and the shaded areas represent standard error of mean. The SPM plots demonstrate the statistical comparison at each frequency and at each contraction level bin between two sides during both index finger abduction (f–j) in the upper panel and elbow flexion (f–j) in the lower panel. The solid red lines indicate the critical value with the significance level set at 0.05 and the dashed red lines the critical value with the significance level set at 0.1, beyond which is the frequence range with significant differences between two sides.

### E. Heterogeneity of muscle impairment between FDI and elbow flexors

The MVIC ratio and FSD ratio demonstrate a measurable heterogeneity of reductions in muscle strength and in force variability (Fig. 7). In terms of muscle weakness, although the average MVIC ratio was comparable between the sides (Table 1), a higher degree of weakness of the elbow flexors was observed in five subjects (i.e., S01, S02, S03, S08, and S10), a higher degree of the FDI muscle in two subjects (i.e., S07 and S09), and a comparable degree between the muscle groups in five subjects (i.e., S04, S05, S06, S11, and S12). Interestingly, the extent to which force variability was affected between the muscle groups was different compared to the impairment in muscle weakness. In particular, four subjects (i.e., S01, S02, S05, and S08) showed greater impairment in force variability of the elbow flexors, whereas six subjects (i.e., S04, S06, S09, S10, S11, and S12) exhibited greater weakness in the FDI muscle.

**Fig. 7.**
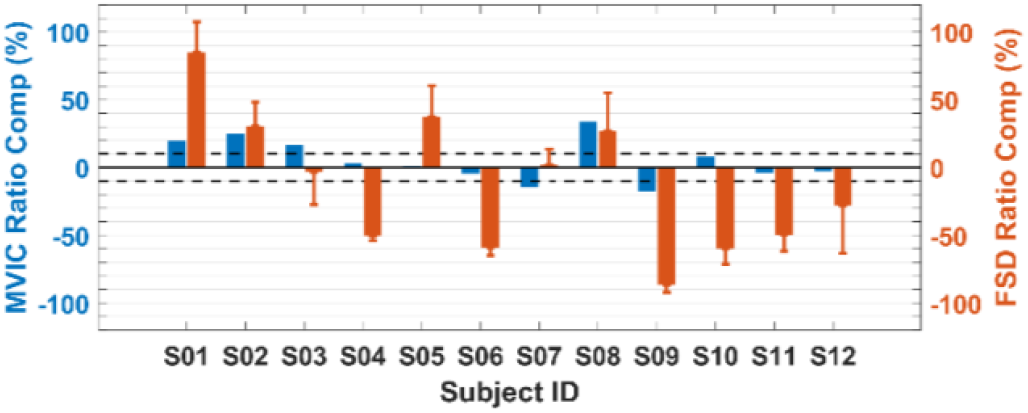
The comparison of relative impairments in muscle weakness and force variability between index finger abduction and elbow flexion for each subject. Error bar indicates standard error of mean across different contraction levels.

### F. Correlation of changes in force variability and muscle weakness

During finger abduction, our results revealed a moderate negative correlation between the increase in force variability and MVIC Ratio at 40% and 50%MVIC, and a weak negative correlation at 60% MVIC, though none were statistically significant. For elbow flexion, the increase in force variability displayed a moderate to strong negative relation with MVIC Ratio, with the significant correlations observed at 20% and 60%MVIC.

## IV. Discussion

This study investigated the changes in force variability during index finger abduction and elbow flexion within the same group of individuals post stroke. Our main findings were: 1) Impairment in force variability is observed on the paretic side across a wide range of force levels during index finger abduction but is not evident until high contraction levels during elbow flexion; 2) The relative increase in force variability on the paretic side compared to the contralateral side is a linear function of contraction level during elbow flexion, but remains constant regardless of contraction level during index finger abduction; 3) A similar frequency distribution in force output is observed on both sides during index finger abduction and elbow flexion, with the peak frequency below 2 Hz, and high-frequency spectral components (i.e., > 2 Hz) in force output appear to increase on the paretic side compared to the contralateral side but only during index finger abduction; 4) Relative changes in the impairment of muscle weakness and force variability are not comparable between the two muscle groups and among the participants. These findings suggest that force variability is generally increased in both FDI and elbow flexors on the paretic side following a stroke, but the degree of such changes may vary by muscle groups and by individuals.

There are several possible mechanisms that contribute to the increased variability of the forces recorded from both FDI or elbow flexors on the paretic side: alterations in MU firing behavior (e.g., firing rate); and changes in neuromechanical properties. If such changes in force variability result from changes in MU firing rate, this could in turn be an outcome of adaptations originating from various locations within the nervous system. For example, it could arise from changes in cortical control, or from changes arising at the spinal level. Variability might be caused by greater dispersion of inter spike intervals, or from the superposition of other periodicities such as tremor or reflex oscillation, including clonus, which is often seen after stroke [16]. We do not yet know which of these factors is important, except to say that there are no obvious or major periodicities visible in the force spectrum during either index finger abduction or elbow flexion. In addition, variations in the degree of changes in force variability across different muscle groups provide insights into the potential modulators of force variability such as muscle fiber composition or neural control pathways for muscle activation.

Greater force variability was consistently observed on the paretic side during index finger abduction, spanning a wide range of contraction levels, except at 20%MVIC (p = 0.057). This observation supports previous findings that individuals with neurological disorders may experience impaired force variability in paretic muscles [7] [8] [9] [10]. For instance, previous studies have highlighted greater force variability in paretic muscles during tasks including wrist extension [10] and power grip [9]. Similarly, the greater force variability (quantified by CoV for force) was observed at low contraction levels in older adults during isometric index finger abduction. It was suggested that this elevation is associated with increased MU discharge variability in older adults [3]. Given that some studies also reported that increased discharge variability in MUs might contribute to motor dysfunction in individuals with mild-to-moderate Parkinson’s disease [27], these findings both shed light on the potential role of MU physiological properties in regulating force variability.

Although there is few evidence on the potential mechanisms underlying the increased force variability in paretic muscles post stoke, it is plausible that such increased force variability could be attributed to abnormalities in MU firing behavior following stroke (e.g., measurable reduction in MU discharge rate [6]). To explain, recordings of many concurrently active MUs from human muscles have indicated that larger MUs recruited later tend to fire at lower rates than those recruited earlier, forming a layered appearance called “onion skin” pattern [4][5]. The firing rate of larger MUs potentially deviates significantly from their optimal fusion frequency, resulting in larger force ripples as force output increases. Consequently, lower discharge rates of MUs post stroke likely led to a relatively unfused muscle contraction and thus larger ripples in force output, manifested as an increase in force variation. Future studies are needed to examine the potential association between force variability and MU firing behavior during isometric contractions.

However, our study revealed that the changes in force variability on the paretic side are less evident during elbow flexion, with an observable increase only at high contraction levels (i.e., 50 and 60%MVIC), indicating that changes in force variability post stroke may vary among muscles. Interestingly, such discrepancy is also noted in older adults. For example, older adults showed a substantial increase in CoV for force during isometric index finger abduction by ∼120% compared to age-matched young individuals [28], while it appears that age has no effects on force variability during isometric elbow flexion [29].

There are several possible explanations for such a different extent of changes in force variability involving elbow flexors. The first one concerns the potential difference in motor unit recruitment strategies and in muscle fiber composition between elbow flexors and FDI. To be precise, the majority of MUs in FDI muscle are recruited from 20%MVIC [30], while the upper limit of the MU recruitment is up to ∼80%MVIC for elbow flexors [23] [31]. This implies that at comparable low force levels, the index finger abduction with FDI involves a larger portion of high-threshold MUs innervated to fast-twitch muscle fibers with greater twitch amplitude and shorter half relaxation time, compared to elbow flexion. It has also been reported that FDI is more of fast-twitch fibers compared to elbow flexors [20] [32]Considering the common post-stroke changes in the MU mechanics (i.e., the lower mean MU firing rate [6] and selective fast-twitch muscle fiber atrophy [21] [33]), the findings that the pronounced differences in FSD between sides were found from low contraction levels in FDI (Fig. 3a) but at high contraction levels in elbow flexors (albeit not significant; Fig. 3b) could be explained in part, which is further manifested in the gradual increase in the relative changes in FSD between sides with increasing elbow flexor force (Fig 4b). Future studies are required to test whether the changes in MU firing rate for these muscles are comparable, which could also contribute to such discrepancies.

The second factor that may contribute to different changes in force variability during finger abduction and elbow flexion is the number of muscles involved in the target tasks. FDI muscle itself can account for most of the abduction force exerted by the index finger, however, multiple muscles are coordinated during elbow flexion. For stroke survivors who often experience weakness in multiple upper extremity muscles, weaker muscles can be compensated by other agonists which are less impaired, therefore, negligible changes in net force variability on the paretic side are more likely. This finding suggests that care should be taken when considering joint force variability as a biomarker for assessing motor impairment in a specific muscle. In addition, different neural control mechanisms might also impact the varying extent of changes in force variability observed in FDI and elbow flexors after a stroke. Unlike FDI with more direct corticospinal projections, elbow flexors primarily rely on projections through spinal interneurons. Moreover, there are alternative descending motor pathways that can regulate the arm function, such as the reticulospinal tract [34]. These alternative descending pathways can partially compensate for deficits in the corticospinal tract. Therefore, the force control capacity post-stroke during elbow flexion might be less susceptible compared to hand muscles, potentially leading to less evident changes in force variability. Further research is needed to explore the potential associations between force variability and neural pathways after a stroke.

Other factors may also contribute to force variability. For instance, age-related physiological and functional changes have been shown to noticeably increase force variability [28] [29]. Further analyses using our dataset revealed no significant effects of age on force variability on both sides, indicating that age may not be a dominant factor to the increased force variability in our samples. However, considering the skewed distribution of age in our samples and the limited sample size, further investigation involving subjects across a broader age range is necessary to examine potential interactions between age and force control impairment after stroke. Additionally, muscle mechanical properties can be substantially altered in individuals with stroke (e.g., altered muscle-tendon unit properties [35] [36] and increased muscle stiffness [37]). Previous studies using mechanomyogram as an alternative measure of force variability, have shown that the oscillation amplitude during isometric contractions decreases with increasing muscle length and muscle stiffness [38]. It would be valuable to determine the relative contributions of stroke-related neural and mechanical alterations to force variability. Visuomotor corrections and feedback gain can also significantly influence force variability during a sustained isometric contraction [39] [40] [41]. Stroke often leads broadly to impairments in sensory, motor, and cognitive functions [42], and in turn the reduced performance of sensorimotor integration [43]. In addition, it has been suggested that force variability is also significantly influenced by visual gain in control subjects [44], which could explain the measurable increase in force variability in FDI compared to elbow flexors (note that the absolute MVIC values of FDI are smaller than those of elbow flexors). Although we compared the force variability parameters at a matched force between sides, the visual gain was not fully controlled across sides, muscles, and participants. It would be interesting to study the potential contributions of these factors to force variability in stroke survivors.

The analysis of the force spectrum reveals that most of the power of the force fluctuation signals were located below 2 Hz, as also reported in the previous studies [45]. Moreover, the paretic FDI muscles demonstrated an increase in the power at ∼0.5 Hz, but there was no significant difference in the power during elbow flexion. Considering the fact that the paretic side showed a greater power at ∼0.2 Hz but smaller at ∼0.6 Hz during grip tasks post stroke [9], the characteristics of force variability as well as task-dependent muscle coordination might be reflected in the force spectrum. There are possible mechanisms that could explain such a low-frequency component in the force fluctuation during a sustained isometric contraction. For instance, low-frequency oscillations in force might be related to harmonic frequencies of the resonant frequency of contracting muscle(s) modeled as a vibrating beam [46]. Moreover, the respiratory frequency (typically falling within the range of 0.2–0.3 Hz, but stroke could alter the respiratory capacity and rhythm [47]) appears influence force fluctuations.

Interestingly, our data demonstrated the trend of a secondary peak beyond 2 Hz on the paretic side during index finger flexion with FDI (Fig. 6 & 7). Although underlying mechanisms are not clear yet, the selective loss of fast-twitch motor units on the paretic side could lead to a compensatory increase in the firing rate of the remaining slow-twitch fibers. This phenomenon might be reflected in the force spectrum. As discussed above, it is plausible that the force fluctuation could stem from adverse effects of “onion skin” firing protocols. However, there exists a disparity between the median frequency of the force spectrum and the discharge rate of individual MUs which is typically greater than 5 Hz [4]. Aforementioned factors related to visuomotor corrections and feedback/visual gains could also result in a higher-frequency component in force output. Further research is required to investigate the underlying sources of force spectrum during a sustained isometric force production and to understand its association with stroke-related changes in force variability.

The present study firstly provided evidence that the degree of motor impairments (i.e., estimated by changes in muscle weakness and in force variability after stroke) may not be uniform across muscles on the same affected side post stroke (Fig. 5). This high heterogeneity could be attributed to many factors including CNS lesion location, different control pathways, and lifestyles (i.e., less use of the paretic hand in daily life) [34] [48] [49]. Furthermore, our results imply that the degree of muscle weakness does not always correlate with the level of impairment in force variability either within a specific muscle or when comparing different muscles. Further research is necessary to investigate the potential dominant causes of such heterogeneity and to assess the effectiveness of muscle weakness and/or force variability measures in quantifying motor impairments.

The correlation analysis between force variability and MVIC Ratio implies that individuals with greater muscle weakness tend to exhibit higher force variability following stroke. Although the linear relation is not always statistically significant (Table 2), the majority of participants with reduced muscle strength consistently display increased force variability. This suggests that tracking force variability could be a valuable biomarker for assessing motor control impairments in clinical populations. However, further studies with larger sample sizes are necessary to validate its consistency and reliability as a diagnostic tool in clinical settings.

**TABLE 2.** Correlation Performance Between fsd_NORM_ AND MVIC Ratio.

|  | Force Level (%MVIC) |  |  |  |  |
| --- | --- | --- | --- | --- | --- |
|  | 20 | 30 | 40 | 50 | 60 |
| Index Finger Abduction | 0.22<br>(0.50) | -0.05<br>(0.88) | -0.43<br>(0.16) | -0.68<br>(0.05) | -0.18<br>(0.61) |
| Elbow Flexion | **-0.74\***<br>(0.01) | -0.31<br>(0.37) | -0.45<br>(0.15) | -0.48<br>(0.19) | **-0.83\***<br>(0.003) |

Note: Data are shown as correlation coefficient $r(p)$ . \* $p < 0.05$ .

There are several limitations in this study. First, our study was cross-sectional in nature and focused on the population of stroke survivors with mild to moderate impairment; thus, caution is required when extrapolating these findings to longitudinal changes in force variability or to individuals with moderate to severe impairment. Moreover, age-matched controls were not included in this study, and the contralateral side was used as a control. The function on the side ipsilateral to the cerebral lesion could also be disturbed [50] [51], although the contralateral limbs in our samples did not show any detectable paresis or spasticity. Lastly, optimal joint positions may vary between sides and among participants, likely due to post-stroke alterations in muscle-tendon unit properties [35] [36] and muscle synergy [52], which were not assessed in this study.

## V. Conclusion

This study suggested that force variability is typically impaired during finger abduction and elbow flexion following a stroke, with the degree of the impairment differing between muscle groups and among participants. Such differences appear to be reflected in the varied dependence of changes in force variability on contraction level as well as in the force spectrum. Correlations between force variability and muscle weakness suggest that force variability could a useful metric for assessing motor control impairment in clinical settings. Future studies are required to explore how post-stroke neuromuscular adaptations can impact force variability and to investigate the relative contribution of such adaptations to heterogeneous motor impairments.

## VI. Acknowledgement

We thank all participants in this study. We also thank Alexander Barry for technical support.

