## Supplemental Figure 1 for "Force variability is a potential biomarker of motor impairment in hemispheric stroke survivors"

**Fig. S1.** Individual Plots for force variability against force level on both sides for each subjects.

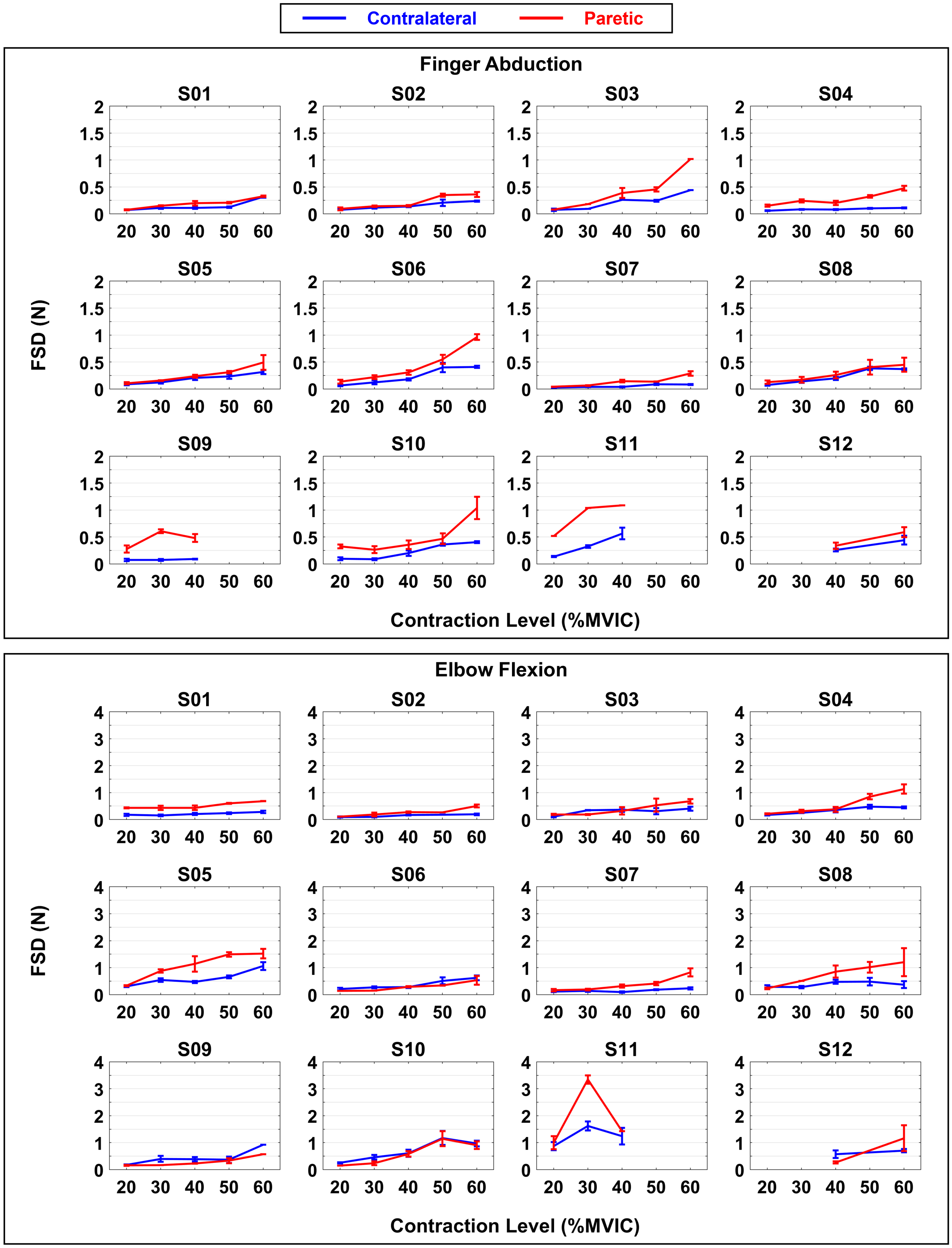
